# Strong heterologous electron sink outcompetes alternative electron transport pathways in photosynthesis

**DOI:** 10.1101/2024.03.18.585510

**Authors:** Michal Hubáček, Laura T. Wey, Robert Kourist, Lenny Malihan-Yap, Lauri Nikkanen, Yagut Allahverdiyeva

## Abstract

Improvement of photosynthesis requires a thorough understanding of electron partitioning under both natural and strong electron sink conditions. We applied a wide array of state-of-the-art biophysical and biochemical techniques to thoroughly investigate the fate of photosynthetic electrons in the engineered cyanobacterium *Synechocystis* sp. PCC 6803, a blueprint for photosynthetic biotechnology, expressing the heterologous gene for ene-reductase, YqjM. This recombinant enzyme catalyses the reduction of an exogenously added substrate into the desired product by utilising photosynthetically produced NAD(P)H, enabling whole-cell biotransformation. Through coupling the biotransformation reaction with biophysical measurements, we demonstrated that the strong artificial electron sink, outcompetes the natural electron valves, the flavodiiron protein-driven Mehler-like reaction, and cyclic electron transport. These results show that ferredoxin-NAD(P)H-oxidoreductase (FNR) is the preferred route for delivering photosynthetic electrons from reduced ferredoxin and the cellular NADPH/NADP+ ratio as a key factor in orchestrating photosynthetic electron flux. These insights are crucial for understanding molecular mechanisms of photosynthetic electron transport and harnessing photosynthesis for sustainable bioproduction by engineering the cellular source/sink balance. Furthermore, we conclude that identifying the bioenergetic bottleneck of a heterologous electron sink is a crucial prerequisite for targeted engineering of photosynthetic biotransformation platforms.

**Significance statement:** We coupled the photosynthetic and biocatalytic (whole-cell biotransformation) performance of model cyanobacteria. We employed a heterologous NAD(P)H utilising enzyme, as a strong artificial electron sink, allowing us to gain a comprehensive understanding of photosynthetic electron partitioning. We demonstrated that the strong electron sink outcompetes natural electron sinks and cyclic electron transport.

## Introduction

Oxygenic photosynthesis is driven by the photo-oxidation of water, liberating electrons which are eventually used to produce the photosynthetic reductant NADPH for carbon fixation and metabolism (Fig. 1a) (Hitchcock et al., 2022; Mullineaux, 2014; Nikkanen et al., 2021). The soluble electron carrier protein ferredoxin (Fd) is the electron distribution hub of photosynthesis (Fig. 1b) (Nikkanen et al., 2021; Hanke and Mulo, 2013; Goss and Hanke, 2014). However, the factors determining the fate of electrons from Fd remain poorly understood. Elucidating this is imperative towards understanding how photosynthetic organisms balance their light reactions with carbon fixation and downstream cell metabolism in response to changing environmental conditions and to rationally harness photosynthetic microorganisms as green cell factories for sustainable bioproduction (Allahverdiyeva et al., 2021; Barbosa et al., 2023; Hitchcock et al., 2022; Santos-Merino et al., 2023; Tan et al., 2022).

Reduced Fd (Fd_red_) is predominantly oxidised by ferredoxin-NADP^+^-oxidoreductase (FNR) to produce NADPH in the final step of linear electron transport (LET). The generated trans-thylakoidal proton motive force (*pmf*), composed of proton and electrical gradients, is utilised by the ATP synthase to produce ATP. Both NADPH and ATP fuel the Calvin-Benson-Bassham (CBB) cycle and various downstream metabolic processes. However, the LET alone does not generate sufficient ATP to meet the demands of the CBB cycle (Allen, 2002; Kramer et al., 2004). Consequently, auxiliary electron transport (AET) plays a critical role in meeting the metabolic ATP requirements, and regulating and protecting the photosynthetic apparatus, particularly under fluctuating environmental conditions. In cyclic electron transport (CET), Fd_red_ is oxidised either by the NDH-1 complex, also known as photosynthetic complex I, in several plant taxa and cyanobacteria, or by the pathway dependent on the PGRL1 and PGR5 proteins in plants and algae (Buchert et al., 2020; DalCorso et al., 2008; Nikkanen et al., 2020; Peltier et al., 2016; Yadav et al., 2020).

Fd_red_ has been recently pinpointed as the electron donor for flavodiiron proteins (FDPs) (Nikkanen et al., 2023, 2020; Sétif et al., 2020), which catalyse the photo-reduction of O_2_ to water (Mehler-like reaction) upon sudden increases in light intensity in cyanobacteria, green algae and gymnosperms (Alboresi et al., 2019; Allahverdiyeva et al., 2013; Bag et al., 2022; Santana-Sanchez et al., 2019; Zhang et al., 2012). Fd_red_ also drives the assimilation of metabolites, such as nitrate and nitrite (Flores et al., 2005), and to a lesser extent, sulfite (Kaneko et al., 1996), glutamate (Navarro et al., 2000) and biliverdin (Frankenberg et al., 2001). Under specific conditions, some green algae and cyanobacteria employ hydrogenases to utilise Fd_red_ for hydrogen metabolism (Kosourov et al., 2021) and N_2_-fixing cyanobacteria use Fd_red_ as the electron donor to nitrogenase (Magnuson and Cardona, 2016). Fd_red_ also provides reducing power for the thioredoxin system, which plays a crucial role in regulating various processes, including light-dependent activation of CBB enzymes (Mallén-Ponce et al., 2022). Finally, Fd has also been proposed as a potential electron exit point for photosynthetic microbes during exoelectrogenesis (Wey et al., 2019).

Photosynthetic reducing equivalents, Fd_red_ and NAD(P)H, can be harnessed as electron donors for heterologously expressed oxidoreductases to convert substrates into industrially relevant compounds through whole-cell photo-biotransformation (Jensen et al., 2011; Malihan-Yap et al., 2022; Schmermund et al., 2019). For example, NAD(P)H-dependent enzymes have been exploited for the oxyfunctionalisation of ketones into lactones (Erdem et al., 2022; Tüllinghoff et al., 2022) and for the reduction of maleimides (Assil-Companioni et al., 2020; Barone et al., 2023). Fd_red_ has supplied electrons to cytochrome P450s (Cyt P450), which offer a wide range of reactions (Agustinus and Gillam, 2023; Wlodarczyk et al., 2016). The effect of introducing these artificial electron sinks on the photosynthetic electron transport chain (PETC) has been partially explored. A recent study demonstrated that the heterologous expression of Cyt P450 in *Synechococcus elongatus* PCC 7942 led to enhancements in the quantum yield of photosystem II (PSII) and maximal oxidation of photosystem I (PSI) reaction centre chlorophylls (P700). Furthermore, a co-expression of carbon and electron sinks had a cumulative positive effect on photosynthetic yield (Santos-Merino et al., 2021).

Deleting natural electron sinks has gained popularity as an approach to enhance the yields of whole-cell biotransformation. However, the effects of such modifications on the PETC vary. A study using Cyt P450 showed an increase in electron transport through PSII by deleting a subunit of the NDH-1 complex, with no effects on PSI (Berepiki et al., 2018). Contrastingly, deleting the thylakoid-localised respiratory terminal oxidase, aa3-type cytochrome c oxidase (COX), led to an increase in electron transport mainly through PSI with a minor increase through PSII (Torrado et al., 2022). Deleting FDPs has also led to an increased biotransformation yield, with no or unclear effects on the PETC (Assil-Companioni et al., 2020; Erdem et al., 2022; Spasic et al., 2022). Such deletions improved the yield often only under specific conditions, indicating the complexity of cellular redox balance and suggesting that the improvements might not be a direct effect of rewiring the electron flux (Assil-Companioni et al., 2020; Santos-Merino et al., 2021; Spasic et al., 2022). A holistic overview of the photosynthetic apparatus and the dynamics of interaction between natural and artificial electron sinks is still missing.

Our primary aim was to elucidate the electron partitioning downstream of PSI and evaluate the effect of a strong artificial electron sink coupled to the photosynthetic light reactions. To achieve this, we combined analytical methods for monitoring the biotransformation reaction with real-time monitoring of O_2_ gas fluxes for the first time. We used previously engineered cyanobacterium *Synechocystis* sp. PCC 6803 (hereafter *Synechocystis*), which heterologously expressed the gene for NAD(P)H-dependent ene-reductase YqjM, sourced from the non-photosynthetic bacterium *Bacillus subtilis* (Fitzpatrick et al., 2003) (Fig. 1c). This recombinant enzyme catalyses the reduction of exogenously added 2-methylmaleimide (2-MM) into 2-methylsuccinimide (2-MS), enabling whole-cell biotransformation. This system has demonstrated fast conversion of 10 mM 2-MM within an hour (Assil-Companioni et al., 2020) and is presumed NAD(P)H-limited under photoautotrophic conditions (Barone et al., 2023).

We carried out a comprehensive analysis of the photosynthetic performance of YqjM-expressing cyanobacteria using a suite of state-of-the-art biophysical and biochemical techniques. This approach allowed us to reveal that the heterologous electron sink YqjM outcompetes the endogenous auxiliary electron transport pathways downstream of PSI for photosynthetic reductants. This indicates that FNR is the preferred route of electrons from the Fd hub if FNR’s activity is not limited by the availability of NADP^+^. Furthermore, YqjM consumed over half of the available photosynthetic electrons for the reduction of 2-MM and effectively prevented the transient pooling of electrons in the PETC. In addition to elucidating the fundamental aspects of the orchestration of photosynthetic electron flux, our findings lay the foundations for rational optimisation of whole-cell biotransformations and other applications that harness the reducing power of photosynthesis.

**Figure 1.**
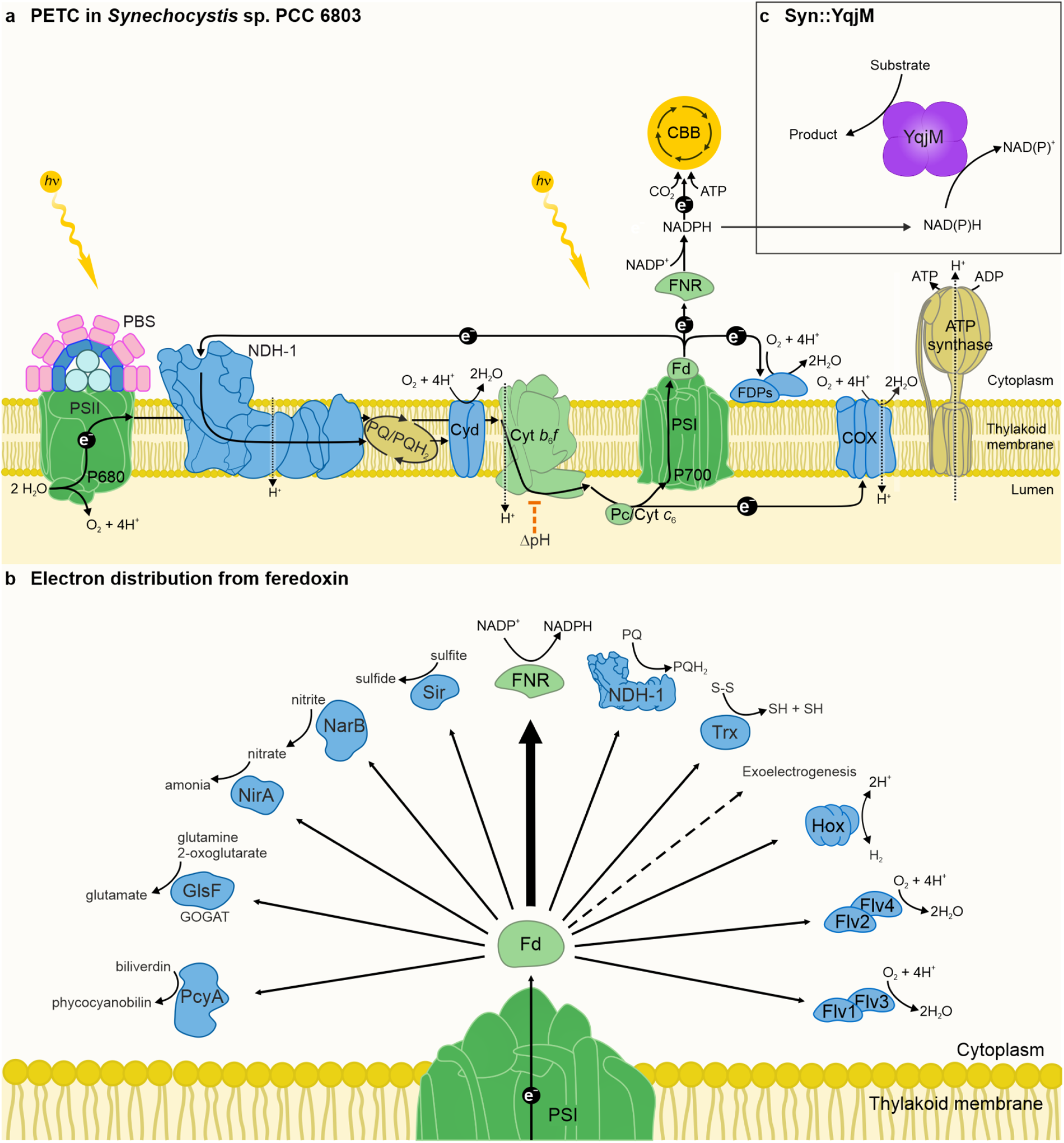
Photosynthetic electron transport chain (PETC) and electron distribution from Fd in the model cyanobacterium Synechocystis sp. PCC 6803. a: Simplified schematic of the PETC. The PETC in cyanobacteria consists of photosystem I and II (PSI and PSII, respectively), Cytochrome b6f (Cyt b6f), and electron carriers plastocyanin (Pc), Cytochrome c6 (present under copper limitation) and ferredoxin (Fd). The last step of linear electron transport (LET) is the reduction of NADP^+^ into NADPH by ferredoxin-NAD(P)H-oxidoreductase (FNR). NADPH is consumed by the Calvin-Benson-Bassham (CBB) cycle and other metabolic processes. Generated pmf is driving ATP synthesis. The cyclic electron transport (CET) around PSI is mediated by the NADH-dehydrogenase-like complex (NDH-1). Respiratory terminal oxidases (RTOs) cytochrome bd quinol oxidase (Cyd) and aa3-type cytochrome c oxidase (COX) reduce O2 to water. Furthermore, flavodiiron proteins (FDPs), constituting a key photoprotective mechanism under fluctuating light conditions, have been lost in angiosperms and red and brown algae (Allahverdiyeva et al., 2015; Ilík et al., 2017; Peltier et al., 2010). b: The electron distribution from Fd. Fdred serves as the electron donor for several pathways. The FDPs in heterooligomeric conformations Flv1/3 and Flv2/4 perform the Mehler-like reaction, reducing O2 to water. The CET, mediated by NDH-1, shuffles electrons from Fd back to the PQ pool. Bidirectional hydrogenase (Hox) is involved in hydrogen metabolism. The thioredoxin (Trx) regulation system regulates, e.g. light-dependent activation of CBB enzymes. Other pathways include phycocyanobilin:ferredoxin oxidoreductase (PcyA), ferredoxin-dependent glutamate synthase (GlsF, GOGAT), ferredoxin-sulfite reductase (Sir), nitrate reductase (NarB), and nitrite reductase (NirA). c: Synechocystis sp. PCC 6803 expressing the gene of the NAD(P)H-accepting flavin-dependent oxidoreductase YqjM (Syn::YqjM).

## Materials and Methods

### Strains and culturing conditions

*Synechocystis* sp. PCC 6803 was used as the WT control in this study. *Synechocystis* strain harbouring the gene coding for the YqjM enzyme under the control of *cpcB* promoter and the substrate 2-MM were used as described previously (Assil-Companioni et al., 2020). Photoautotrophic cultivation was performed in 30 mL of BG-11 medium (20 mM HEPES, pH = 7.5) in 100 mL Erlenmeyer flasks. The cultivation conditions were 30 °C, ambient CO_2_ (0.04%), 50 µmol_photons_ m^-2^ s^-1^ constant white illumination and orbital shaking at 115 rpm. Kanamycin (50 µg mL^-1^) was used in pre-experimental and stock cultures. No antibiotic was present in the experimental cultures.

### Biotransformation conditions

Before biotransformation, cells were harvested in the logarithmic growth phase by centrifugation (5000 *g*, 8 min, room temperature) and adjusted to OD_750_ = 2.5 in fresh BG-11. Chlorophyll *a* (Chl *a*) concentration was quantified by extraction with 90% methanol. The extract was measured at OD_665_ and multiplied by 12.7 to obtain the final concentration in mg mL^-1^. The biotransformation conditions were adapted from a previous study (Assil-Companioni et al., 2020), with these adjustments: illumination of 200 µmol_photons_ m^-2^ s^-1^ white light, 2-MM concentration 10 mM and five minutes of reaction time. After the five minutes biotransformation reaction, samples were taken for specific measurements described below. To calculate the specific activity of YqjM, the biotransformation was followed for 3 hours with samples taken periodically, rapidly frozen in liquid nitrogen and stored at - 80 °C prior to gas chromatography (GC) analysis.

### Membrane Inlet Mass Spectrometry (MIMS)

The *in vivo* fluxes of ^16^O_2_ (m/z = 32), ^18^O_2_ (m/z = 36), and CO_2_ (m/z = 44) were followed by built-in-house MIMS as described earlier (Mustila et al., 2016) with minor adjustments. The final Chl *a* concentration was ∼6 µg mL^-1^, the total dissolved inorganic carbon concentration was adjusted to 1.5 mM by adding NaHCO_3_, and the illumination period was set to 5-5-5 minutes of dark-light-dark, respectively. During the sample preparation, samples were diluted to the appropriate Chl *a* concentration with fresh BG-11 and handled in darkness. The gas exchange rates were calculated as reported (Beckmann et al., 2009). The rates are calculated from a 30s sliding window, thus they appear to increase prior to the onset of illumination and decrease before the end of the illumination period.

### Fluorescence and absorbance changes

Chl *a* fluorescence was followed with the DUAL-PAM 100 spectrophotometer (Walz, Effeltrich, Germany). After the biotransformation reaction, samples were dark adapted for two minutes before being exposed to 100 µmol_photons_ m^-2^ s^-1^ red actinic light (AL) for 240 s, followed by another 240 s of 170 µmol_photons_ m^-2^ s^-1^ red AL illumination. An additional 60 s were recorded with no AL illumination to follow the dark relaxation of the fluorescence signal to observe the F_0_ rise.

DUAL-KLAS-NIR (DKN) spectrophotometer (Walz, Effeltrich, Germany) was used to follow the redox changes of Pc, P700 and Fd as described previously (Nikkanen et al., 2020). Briefly, absorbance changes between 780-820, 820-870, 840-965 and 870-965 nm were used to deconvolute the redox changes of Pc, P700 and Fd using differential model plots. A set of *Synechocystis* mutant strains was previously used to measure these differential model plots using scripts provided by the manufacturer (Nikkanen et al., 2020). Before the measurement, Chl *a* concentration was set to 10 µg mL^-1^ by adding fresh BG-11, and samples were dark adapted for 10 minutes. The redox changes were followed by the modified NIRMAX script supplied in the software package, which is made up of 3 s red AL illumination (3400 µmol_photons_ m^-2^ s^-1^) followed by 4 s of the dark period and an additional 10 s of far red-light illumination. A saturating pulse to fully reduce the Fd pool was introduced after 200 ms of red AL illumination. Due to copper-replete BG-11 medium, Pc is the dominant electron carrier from Cyt *b_6_f* to PSI. Any Cyt *c_6_* present could have a minor effect on the Pc signal (Sétif et al., 2019).

NAD(P)H fluorescence was measured with the NADPH/9-AA module for DUAL-PAM-100 (Walz, Germany). Samples were diluted with BG-11 to reach Chl *a* concentration of 5 µg mL^-1^ and dark-adapted for 10 min. The light-induced NADPH accumulation was followed by 10 s of dark period followed by 180 s of red AL illumination (200 μmol m^-2^ s^-1^) and a minute-long dark period.

### Redox changes of Cyt *f* and *b* hemes

The redox changes of Cyt *f* and *b* hemes of the Cyt b_6_*f* complex were deconvoluted from the absorbance changes at 546, 554, 563 and 573 nm measured with the JTS-10 spectrophotometer (BioLogic, Seyssinet-Pariset, France), appropriate 10 nm full width at half-maximal (FWHM) interference filters (EdmundOptics, Barrington, NJ, USA) and BG39 filters (Schott, Mainz, Germany) protecting the light detectors from scattering effects. The samples were adjusted to Chl *a* concentration of 5 µg mL^-1^, and dark-adapted for three minutes prior to measurement with each filter. Then, they were illuminated for 5 s with 500 µmol_photons_ m^-2^ s^-1^ green AL, and white detection flashes were administered during 200 µs dark intervals.

### Cyclic electron transport ratio quantification

Linear and cyclic electron transport rates were quantified by measuring dark interval relaxation kinetics (DIRK) of the P700 and Pc signals using the DKN, as described previously (Theune et al., 2021). The samples were adjusted to Chl *a* concentration of 15 µg mL^-1^, Briefly, cells were pre-illuminated for 2 min under 500 µmol_photons_ m^-2^ s^-1^, after which the light was repeatedly (100x) shut off for 20 ms. The signals were averaged, and the electron flow through PSI was calculated. Samples were measured in the presence and absence of 20 µM DCMU.

### Calculating *pmf* from electrochromic shift (ECS) measurement

The ECS was measured as described previously (Nikkanen et al., 2020). Briefly, samples were set to a final Chl *a* concentration of 7.5 µg mL^-1^ and dark-adapted prior to exposure to 500 µmol_photons_ m^-2^ s^-1^ green AL interrupted with dark intervals. *pmf* was calculated as the extent of ECS decrease at the dark intervals. Thylakoid conductivity (gH^+^) was calculated as the inverse of the time constant of a first-order fit to ECS relaxation kinetics during a dark interval, and proton flux (vH^+^) as *pmf* × gH^+^ (Cruz et al., 2005).

### Biotransformation coupled with MIMS

The sample was set in the MIMS sample chamber as described in MIMS methods above with the following modifications. The Chl *a* concentration was set to 11 ± 1 µg mL^-1,^ and the final sample volume was 1 mL. The substrate was added to the final concentration of 10mM, and the measurement was started by 5 min dark-adaptation followed by 10 min of illumination (200 µmol_photons_ m^-2^ s^-1^) and 5 min of a dark period. The reaction was stopped by rapidly freezing the sample in liquid nitrogen, and the samples were stored at -80 °C until GC analysis. The dark rate was calculated from a reaction set up in complete darkness for one hour in otherwise the same conditions.

### Gas chromatography

Samples were extracted using 3-step extraction with ethyl acetate. The organic phase was dried using anhydrous MgSO_4_ and analysed on GC-2010 Pro gas chromatograph (Shimadzu, Japan) equipped with an HP-5MS 30 m × 0.25 mm (5%-Phenyl)-methylpolysiloxane column (19091S-133, Agilent) with nitrogen as the carrier gas with splitless injection mode. Compounds were separated at 35 °C (hold 3 min), 200 °C (hold 3 min, 10 °C min^−1^) and 300 °C (hold 3 min, 25 °C min^−1^). Linear velocity was 11 cm sec^−1^. Acetophenone was used as the internal standard. Calibration was performed with known amounts of 2-MM and 2-MS.

### Protein extraction and immunoblotting

Total protein extracts from cultures were isolated as previously described (Zhang et al., 2009), separated by sodium dodecyl sulfate-polyacrylamide gel electrophoresis on BioRad Mini-PROTEAN TGX 4-15% precast gels with 6 M urea and blotted on polyvinylidene fluoride membranes. Membranes were probed with custom polyclonal antibodies raised against Flv1, Flv2, Flv3 and Flv4 (GenScript, USA). Horseradish peroxidase-conjugated secondary antibody (GE Healthcare, Chicago, IL, USA) and Amersham ECL (GE Healthcare) were used for detection.

### Statistical analysis

Analysis and visualisation were conducted using R statistical software (R Core Team, 2020) and Origin (Version 2016) (“Origin,” 2016). Two-way ANOVA was used to test for significance. Tukey’s test was used to compare the means of different groups with each other, with p<0.05 as the significance cutoff.

### Spectra measurement

The light spectra were measured using SpectraPen (PSI, Czech Republic) under the same conditions used in experiments -500 µmol_photons_ m^-2^ s^-1^. Spectra for MIMS and DKN instruments are shown in Fig. S1.

## Results

### Active heterologous electron sink YqjM is a good tool for studying electron fluxes from Fd

In order to isolate the effects of the active heterologous electron sink on the photosynthetic electron fluxes in cyanobacteria, we confirmed that heterologously expressing YqjM in *Synechocystis* (Syn::YqjM) without the addition of the substrate has no effect on photosynthesis (Fig. S2).

The addition of the substrate 2-MM (substrate, S, 10mM final concentration) (Assil-Companioni et al., 2020; Köninger et al., 2016) to wild-type (WT) cells not expressing YqjM (WT+S) inhibited both gross and net O_2_ evolution. However, in cells expressing YqjM (Syn::YqjM), the addition of the substrate (Syn::YqjM+S), while making the induction of O_2_ evolution faster, did not alter steady-state O_2_ evolution (Fig. 2a, b, Fig. S3a, b). This shows that active YqjM does not perturb the primary electron input into the PETC (water photo-oxidation by PSII). Consequently, it serves as a suitable tool for comparing the impact of active YqjM on photosynthetic electron fluxes in relation to WT. Additionally, the substrate inhibited CO_2_ fixation in both WT and Syn::YqjM cells (Fig. 2c, d, Fig. S4), which allowed us to examine AET centred around Fd (Fig. 1b) isolated from interplay with the carbon metabolism. The addition of substrate to WT cells increased the ratio of O_2_ photoreduction (indicating predominantly FDP activity) relative to gross O_2_ evolution from 48 ± 7% to 81 ± 4% during the dark-to-light transition and from 20 ± 5% to 50 ± 6% at steady-state illumination, while the O_2_ photoreduction rate slightly decreased (Fig. S3c, d, e). Similarly, the CET:LET ratio more than doubled from 14 ± 4% to 35 ± 17% upon the addition of substrate to WT cells (Fig. S5a). The number of electrons directed towards CET also increased four-fold (Fig. S5b). These data indicate relatively greater electron distribution to AET pathways in the presence of the substrate in WT, possibly due to the lack of CO_2_ fixation.

**Figure 2.**
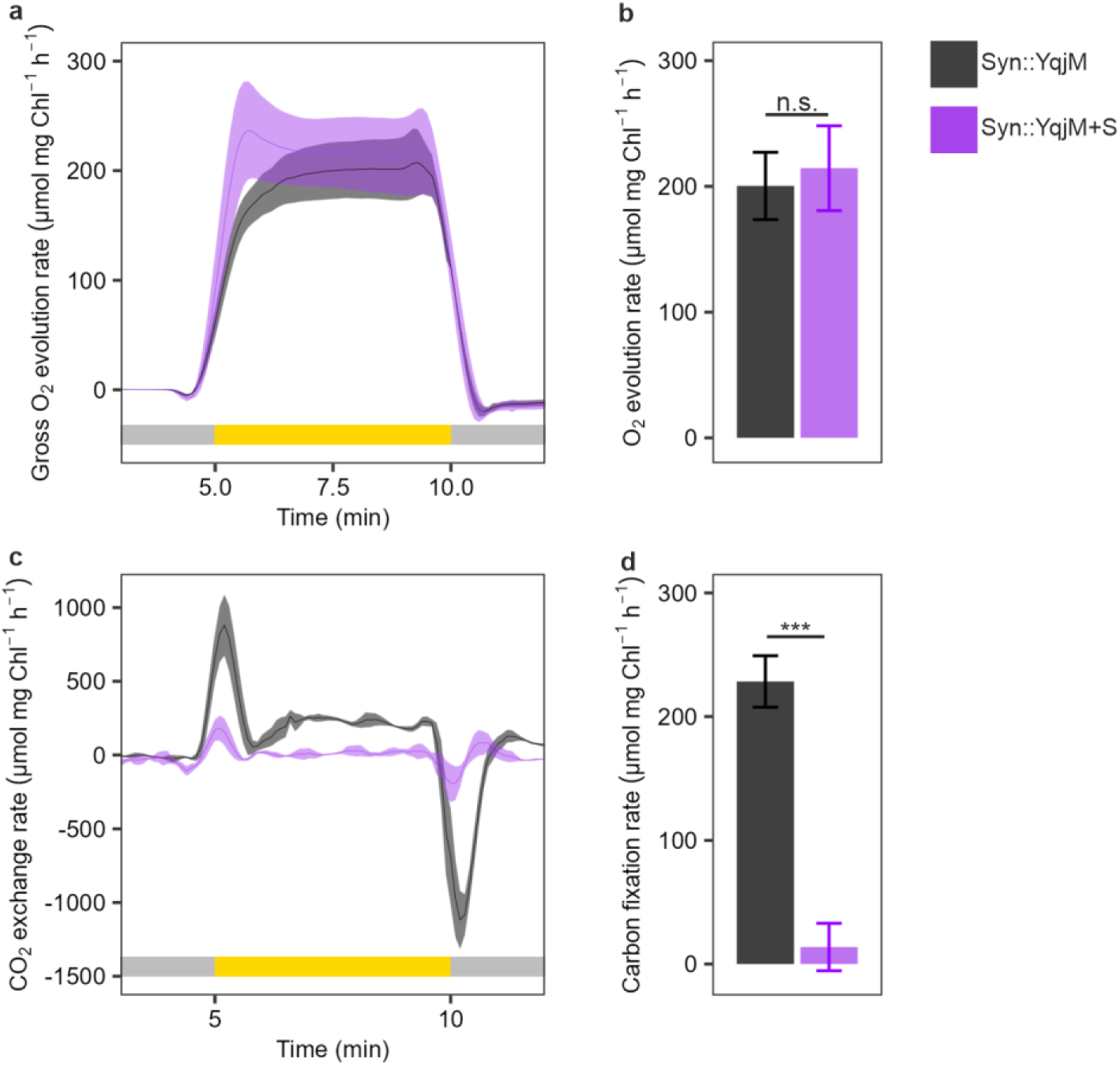
Gas exchange in Syn::YqjM±S. **a:** Kinetics of gross O2 evolution rate. **b:** Steady-state gross (p = 9.05 × 10^−1^) O2 evolution rate. **c:** Kinetics of CO2 exchange rate. **d:** Steady-state carbon fixation rate (p = 1.03 × 10^−4^). Rates are calculated from a 30s sliding window. Data presented as the mean of 3 biological replicates, error bars in column graphs or shading in traces are standard deviations. Statistical significance was tested by one-way ANOVA, n.s. > 0.05, *** ≤ 0.001. Black - Syn::YqjM, purple - Syn::YqjM+S, grey bar in **a** and **c** denotes darkness, yellow bar in **a** and **c** denotes illumination. For gas exchange in WT±S see Fig. S3.

Interestingly, the substrate increases the proton conductivity of the thylakoid membrane in both WT and Syn::YqjM cells (Fig. S6). This results in the dissipation of the *pmf* (Fig. S6), preventing the induction of ‘photosynthetic control’ at Cyt *b_6_f*. The substrate thereby unleashes photosynthesis for uninhibited electron flux to Fd.

### Active heterologous electron sink YqjM outcompetes natural auxiliary electron transport pathways

With active YqjM established as a tool for studying photosynthetic electron fluxes, we probed its effects on the electron distribution downstream of PSI considering Fd_red_ as the distribution hub (Fig. 3a). YqjM converts supplied 2-MM to 2-MS within hours while oxidising NAD(P)H into NAD(P)^+^ (Fig. 1c). Syn::YqjM exhibited specific activity 116 ± 29 U g_DCW_^-1^ [OD_750_ = 2.5, (Fig. 3b)] comparable with previously reported values 166 ± 5 U g_DCW_^-1^ [OD_750_ = 2, (Assil-Companioni et al., 2020)]. It is important to note that while NADH can also serve as the electron donor to YqjM, NADPH has a 3.5-fold higher affinity to YqjM (Assil-Companioni et al., 2020). YqjM maintains only residual activity in the dark, demonstrating a considerably lower reaction rate (Assil-Companioni et al., 2020). Upon illumination, YqjM efficiently utilised NADPH produced by the light reactions, as shown by the negligible light-induced accumulation of NADPH in Syn::YqjM+S (Fig. 3c, Fig. S7). Similarly, the Fd pool remained oxidised during illumination, as revealed by near-infrared absorbance difference measurements (Fig. 3d), indicating efficient and continuous oxidation of Fd_red_.

Next, we investigated if the AET pathways, reliant on Fd_red_ as their electron donor, contribute to Fd oxidation in Syn::YqjM+S. The activity of NDH-1 mediated CET was followed by the post-illumination increase in chlorophyll fluorescence (F_0_ rise), and the CET:LET ratio was determined by analysing the dark-interval relaxation kinetics of the P700 and Pc redox change signals using the DUAL-KLAS-NIR (DKN) (Theune et al., 2021). While the F_0_ rise was eliminated, the CET:LET ratio decreased 6-fold from 19 ± 13% to 3 ± 1% in Syn::YqjM+S (Fig. 3e). The O_2_ photoreduction catalysed by FDPs was monitored by Membrane Inlet Mass Spectroscopy (MIMS), using the enrichment of a heavy oxygen isotope (^18^O_2_) during real-time gas exchange measurements to distinguish O_2_ evolution from consumption. With YqjM active and oxidising NAD(P)H, O_2_ photoreduction decreased close to non-detectable values (Fig. 3f). As a result, the ratio of O_2_ photoreduction to gross O_2_ evolution at the dark-to-light transition and steady-state illumination also strongly decreased (Fig. S3d, e). Importantly, FDP accumulation levels were unchanged in Syn::YqjM compared to WT (Fig. S8). These results suggest that when the NADPH/NADP^+^ pool is kept oxidised by active YqjM in Syn::YqjM+S, the available Fd_red_ is primarily used by FNR for NADP^+^ reduction.

**Figure 3.**
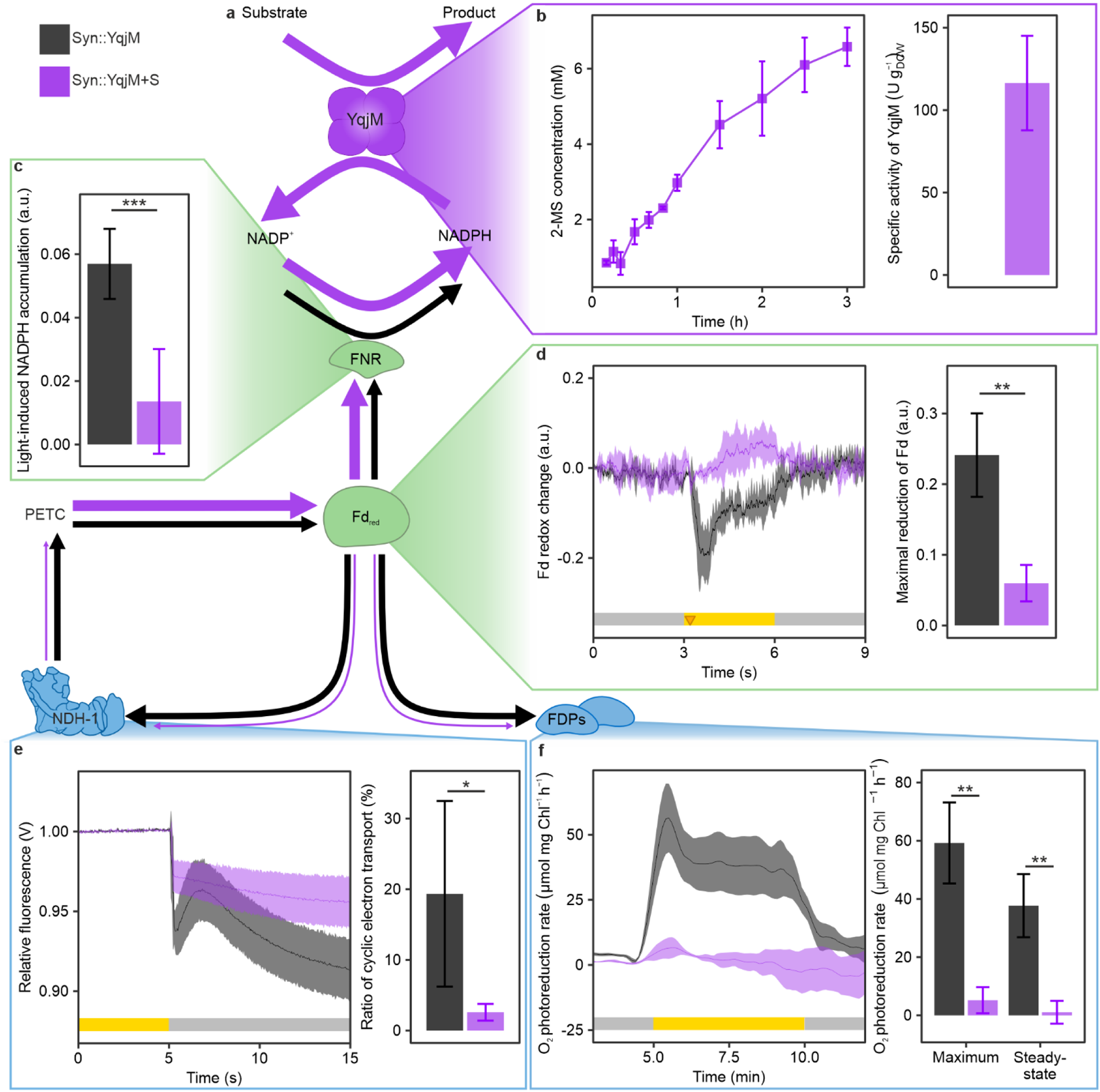
The electron distribution downstream of PSI in Syn::YqjM±S. **a:** Schematic of electron flux. The black arrows signify Syn::YqjM, while the purple arrows signify Syn::YqjM+S. The arrows’ thickness indicates the electron flow. **b:** Concentration of the product during the biotransformation reaction and specific activity of YqjM calculated from the first 20 minutes of biotransformation reaction. **c:** Light-induced accumulation of NADPH as the difference between NAD(P)H fluorescence level after and before illumination (p = 6.85 × 10^−4^). **d:** The kinetics of Fd redox state upon illumination and the maximal reduction of Fd upon illumination relative to the dark level (p = 4.43 × 10^−3^). **e:** The post-illumination F0 rise in chlorophyll fluorescence normalised to steady-state fluorescence, and the CET ratio (p = 4.40 × 10^−2^) assessed by dark-interval relaxation kinetics of P700 and Pc during illumination. **f:** The kinetics of O2 photoreduction rate upon illumination, and the maximal (p = 1.40 × 10^−3^) and steady-state (p = 3.58 × 10^−3^) O2 photoreduction rate as a proxy for the activity of FDPs. Rates are calculated from a 30s sliding window. Data presented as the mean of 3 biological replicates (4 in **c** and CET ratio in **e**), error bars in column graphs or shading in traces are standard deviations (standard error of the mean in **b** 2-MS concentration over time). Black - Syn::YqjM, purple - Syn::YqjM+S, the grey bar denotes darkness, and the yellow bar denotes illumination. Statistical significance was tested by one-way ANOVA, * ≤ 0.05, ** ≤ 0.01, *** ≤ 0.001.

The presence of the active YqjM as an electron sink in Syn::YqjM+S approximately doubles the PSI electron transport rate despite the drop in CET contribution. Thus, enhanced LET is the cause of this increase (Fig. 4b). The strong oxidising effect of active YqjM on the PETC was further demonstrated by the absence of transient re-reduction of P700, the Cyt *b_6_f* and the electron carrier Pc during the first seconds of illumination in Syn::YqjM+S (Fig. 4c-f - dashed vertical line). As we use copper-replete media in this study, Pc is the main electron carrier between Cyt *b_6_f* and PSI. The *b* hemes, Cyt *f* and Pc remained oxidised even after the illumination period (Fig. 4c-e, Fig. S9). The efficient oxidation of the PETC downstream of PSII (Cyt *b_6_f*, Pc and P700) by the active YqjM allows for immediate induction of O_2_ evolution upon illumination (Fig. 2a).

**Figure 4.**
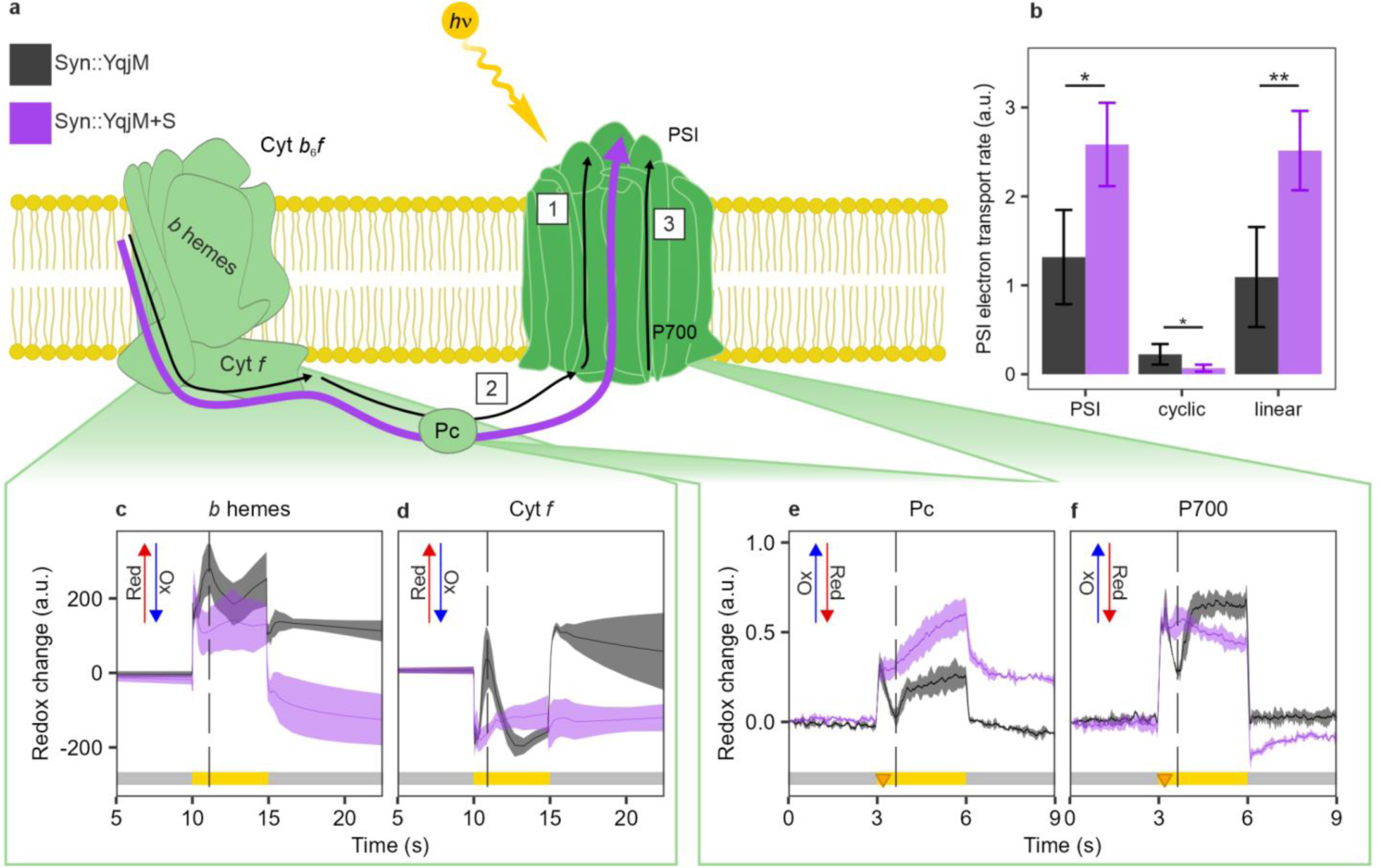
The redox state of the PETC and the electron transport through PSI in Syn::YqjM±S. **a:** Schematic of the oxidation/reduction steps of the PETC upon illumination in Syn::YqjM-S (black) and Syn::YqjM+S (purple). 1. Oxidation of the PETC upon illumination, 2. The transient reduction caused by the inflow of electrons from PSII and the lack of available acceptors downstream of PSI, and 3. The re-oxidation of the PETC when electron acceptors are available. In Syn::YqjM+S, no transient reduction is observed, and the PETC is kept oxidised. **b:** Total electron transport through PSI with WT set to 1 (p = 1.16 × 10^−2^) and the individual cyclic (p = 4.33 × 10^−2^) and linear (p = 7.44 × 10^−3^) electron transport during steady-state illumination. Redox signatures of Cyt b6f components **c:** b hemes and **d:** Cyt f. See also Fig. S10 for the Cyt b6f redox signatures in WT±S. The redox signature of **e:** the electron carrier Pc, and **f:** PSI reaction centre chlorophylls P700. The triangle represents the time a saturating pulse was given to the sample. The dashed vertical line points at the transient reduction of P700, Pc, b hemes and Cyt f. The red arrow signifies reduction (Red), while the blue arrow oxidation (Ox). The coloured bar in **b** - **e** signifies the illumination mode, grey - dark, yellow - illumination. Data presented as the mean of 3 biological replicates (4 in **b**), error bars in column graphs or shading in traces are standard deviations (standard error of the mean in **c and d**). Statistical significance was tested by one-way ANOVA, * ≤ 0.05, ** ≤ 0.01.

### Electron flux analysis: YqjM consumes more than half of the electrons released by PSII

With YqjM efficiently consuming available NADPH produced by the light reactions, we coupled the biotransformation reaction with real-time gas exchange measurements to quantify the flux of electrons from water oxidation to YqjM. After initiating the biotransformation process by introducing 10 mM of substrate to the cells, the reaction was monitored for 20 minutes, out of which 10 minutes were in light and 10 minutes in the dark in the 5-10-5 minutes of dark-light-dark regime (Fig. S11). Subsequently, 0.69 ± 0.07 mM of the product, 2-MS, was detected. By subtracting the dark rate, which was calculated in a separate experiment, we determined that the light-dependent production was 0.45 ± 0.07 mM. Correspondingly, within the 10 minutes illumination period, we measured 0.41 ± 0.03 mM of total evolved O_2_. Hence, the ratio of product formed to O_2_ evolved was 1.09 ± 0.09. PSII oxidises two molecules of water into one molecule of O_2_ through the extraction of four electrons from the water molecules. On the other hand, two electrons are needed to produce each molecule of NADPH, and YqjM uses one molecule of NADPH per one molecule of substrate. We, therefore, concluded that 54 ± 5% of the electrons released by O_2_ evolution at PSII end up being used by YqjM for reduction of the substrate (Fig. 5). As a small portion of the electrons from water oxidation is lost, e.g., as heat due to inefficiencies in the PETC before Fd, YqjM consumes a major portion of the chemical energy available for downstream processes in the form of NADPH.

**Figure 5.**
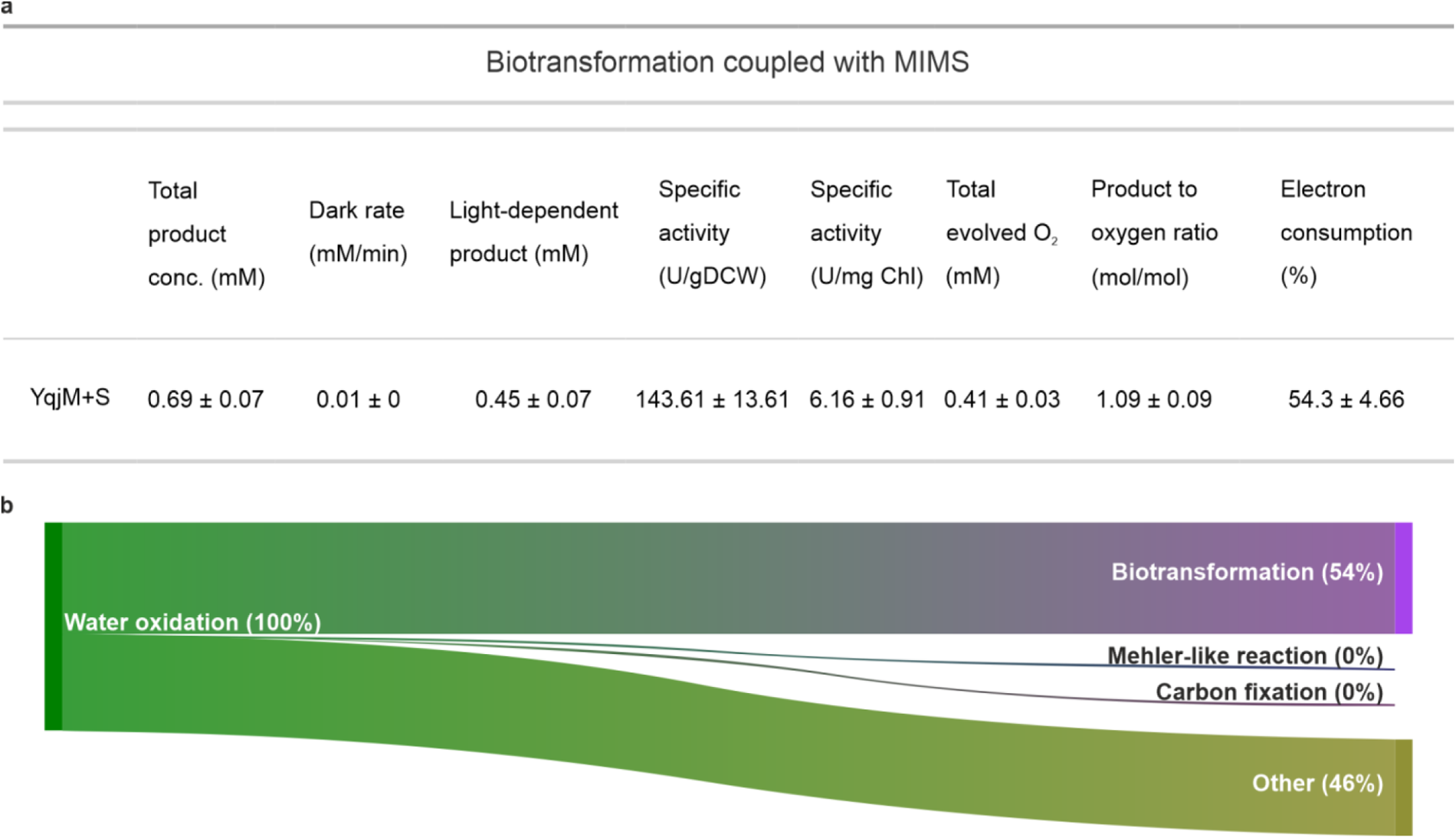
Electron flux towards active YqjM. **a:** Electron flux values derived from three independent biotransformation reactions performed in the MIMS sample chamber measuring the gas exchange followed by product concentration determination by GC. **b:** Mapped electron flux originating from water oxidation towards biotransformation reaction compared to other end-processes.

## Discussion

Orchestrating the distribution of photosynthetic reducing power from Fd is crucial for both photosynthetic organisms responding to change and the bioengineer re-directing photosynthetic reductants to desired reactions. However, the mechanisms determining the fate of electrons at the Fd distribution hub have remained largely elusive. Our results reveal that in the presence of the strong NAD(P)H-consuming heterologous sink, which maintains the NADPH/NADP^+^ pool at an oxidised state (Fig. 3c), the AET pathways of the Mehler-like reaction and CET are outcompeted for electrons from Fd (Fig. 3e, f). This indicates that when the availability of the NADP^+^ substrate is not limiting its activity, FNR is the preferred acceptor for electrons from Fd, pinpointing the cellular NADPH/NADP^+^ ratio as a key factor in orchestrating photosynthetic electron flux. These findings have profound implications for rationally optimising the productivity of biotransformation platforms based on photosynthetic microbes and improving crop plants’ yield and climate resilience.

While incremental advances in understanding how the distribution of photosynthetic reducing power from Fd_red_ is controlled have previously been made, the underlying mechanisms have remained largely unknown. For example, *Synechocystis* has 11 Fd isoforms (Artz et al., 2020), but their specific bioenergetic roles are poorly understood. High-abundance Fd1 is the main isoform involved as the final electron acceptor in the PETC. We recently showed that it is also the main electron donor to FDP hetero-oligomers (Nikkanen et al., 2023). The low-abundance Fd isoforms 2-11 likely serve other condition-specific processes such as ROS scavenging systems (Cassier-Chauvat and Chauvat, 2014; Marteyn et al., 2009), the NiFe-hydrogenase (Gutekunst et al., 2014), nitrogen assimilation (Motomura et al., 2019), or mixotrophic carbon metabolism (Wang et al., 2022). Coordination of electron flux from Fd1 between FNR (and downstream carbon fixation) and the AET pathways is, therefore, the primary task in maintaining maximal photosynthetic efficiency without compromising photoprotection. However, besides the NADPH/NADP^+^ ratio, other regulatory mechanisms likely also play a role in achieving dynamic coordination of electron flux, and several influencing factors must be considered.

We showed that the addition of the 2-MM substrate achieves an uncoupled state of photosynthesis, as the *pmf* is dissipated due to the increased conductivity of the thylakoid membrane (Fig. S6). The dissipation of the *pmf* means that photosynthetic control, i.e. inhibition of PQH_2_ oxidation at Cyt *b_6_f* at low lumenal pH (Malone et al., 2021), is not a limiting factor of YqjM biotransformation yield, and optimisation strategies aiming at *pmf* uncoupling, for example by deletion of the inhibitory epsilon subunit of ATP synthase (Imashimizu et al., 2011) or by using ionophoric uncouplers, will not provide further benefit. Strategies to decrease the CO_2_ fixation activity of CBB as a competitive NADPH sink will be ineffective, as the CBB is already inactive (Fig. 2c, d, Fig. S4), possibly due to disturbed redox regulation caused by the thiol-binding nature of 2-MM. The factor limiting biotransformation yield in Syn::YqjM is likely either water oxidation and consequent LET rate, or the capacity of FNR to reduce NADP^+^ since all NADPH produced by FNR in light was efficiently consumed in the presence of active YqjM (Fig. 3c). FNR activity may also be affected by *pmf* dissipation. In plant chloroplasts, the binding of FNR to its thylakoid anchor proteins is pH-dependent (Benz et al., 2010). While it is yet to be determined if *pmf* also controls the association of FNR to cyanobacterial phycobilisomes, FNR has been observed to disassociate from phycobilisomes in a low ionic strength buffer (Van Thor, 1999).

The association of FDP heterooligomers with the thylakoid membrane increases with the loss of *pmf*, likely enhancing the O_2_ photoreduction activity (Nikkanen et al., 2023). Therefore, adding 2-MM should result in a relative increase in O_2_ photoreduction, which we observed in WT (Fig. S3d, e). In Syn::YqjM, the lack of O_2_ photoreduction in the presence of the substrate indicates that the exogenous NAD(P)H-consuming electron sink is able to outcompete FDPs despite the FDP-activating effect of the *pmf* dissipation. Similarly, the addition of 2-MM increased CET in WT (Fig. S5) but not in Syn::YqjM+S (Fig. 3e, Fig. 4b). This suggests that NDH-1 is outcompeted for electrons from Fd_red_ in the presence of active YqjM. In addition to the above-mentioned effects, 2-MM can potentially have other targets within the PETC (Appendix S1).

Interestingly, LET through PSI strongly increased in the presence of active YqjM, while the O_2_ evolution by PSII showed only a minor, nonsignificant increase (Fig. 4b, Fig. 2a, b). A roughly 30% increase in electron transport rate in PSI with a minor effect on PSII was also observed in a *Synechococcus* strain lacking COX and expressing Cyt P450 (Torrado et al., 2022). Although the inhibition of thylakoid RTOs affected Y(II) and P700 oxidoreduction due to the accumulation of electrons in the PETC, the contribution of RTOs to O_2_ photoreduction in *Synechocystis* is low (Ermakova et al., 2016; Viola et al., 2021). Therefore, it is not likely that the redirection of electrons from Pc towards PSI instead of COX is sufficient to account for the discrepancy in the PSI and PSII electron transport rates. The additional electrons to PSI also do not derive from CET as the presence of active YqjM resulted in the loss of CET (Fig. 3e). It is important to note here that despite PSII and PSI electron transport rates being determined with the same intensity of photosynthetically active radiation (PAR, 400-700nm, 500 µmol_photons_ m^-2^ s^-1^), the actinic light spectra are different (PSII determined by MIMS with broad white, PSI determined by DKN with 623nm red, Fig. S1) resulting in differential excitation of photosystems. Indeed, previous studies have pointed out the challenge of reliable quantification of simultaneous electron transfer rates (Fan et al., 2016; Kauny and Sétif, 2014). Therefore, building coupled systems for simultaneous measurements, as we have achieved with MIMS and biotransformation here (Fig. 5), is an essential approach for achieving a holistic understanding of bioenergetics.

YqjM effectively oxidised available NAD(P)H when supplied with 2-MM substrate. Previously, it has been estimated that YqjM consumes over 60% of NADPH produced by light reactions with a consumption of 680 μmol mg_chla_^−1^ h^−1^ (Assil-Companioni et al., 2020). The estimated NADPH production in *Synechocystis* is 530 - 1070 μmol mg_chla_^−1^ h^−1^ (Kauny and Sétif, 2014), demonstrating that YqjM can utilise most of the NADPH produced by FNR, depending on the specific conditions. Upon isolating the light-dependent production rate, we calculated that YqjM used over half of the electrons originating from water oxidation at PSII to reduce 2-MM into 2-MS (Fig. 5), with the remaining electrons being lost due to inefficiencies in the PETC or consumed in other metabolic pathways downstream of PSI. It is important to note that there is uncertainty in these estimations, and a discrepancy between rates of oxygen evolution and NADPH consumption has been reported (Kauny and Sétif, 2014). Nonetheless, our calculations indicate that a substantial portion of the reducing power produced by the photosynthetic light reactions is consumed by YqjM.

Targeting AET pathways has emerged as a popular strategy for increasing the product yield of whole-cell biotransformation applications (Assil-Companioni et al., 2020; Erdem et al., 2022; Jurkaš et al., 2022). Although the deletion of FDPs led to improved biotransformation yield (Assil-Companioni et al., 2020; Erdem et al., 2022), our results indicate that these enhancements are likely achieved indirectly rather than via the decreased competition for electrons from the PETC. Instead, for rational optimisation, we should focus on increasing the photosynthetic electron transport rate in the PETC, engineering the enzymatic activity, or targeting the localisation of the enzyme. Attaching native and heterologous enzymes to PSI or Fd has been shown to be successful with Cyt P450s (Lassen et al., 2014; Mellor et al., 2016) and hydrogenase (Appel et al., 2020). However, in these applications, Fd_red_ is the electron donor rather than NAD(P)H. A fusion of PSI, Fd and FNR enhanced the electron transport rate (Medipally et al., 2023), demonstrating that the localisation of heterologous enzymes is a promising target for optimisation. Another potential target is the NADPH/NADP^+^ pool. Increasing its overall size by, e.g. enhancing the phosphorylation of NAD^+^ by overexpressing NAD kinases (Ishikawa and Kawai-Yamada, 2019), could supply FNR with additional NADP^+^, allowing it to utilise more Fd_red_. Additionally, considering the effects of the substrate on the cell is of utmost importance when engineering a production strain for whole-cell biotransformation.

In conclusion, we demonstrate that the strong artificial electron sink outcompetes the natural electron valve, flavodiiron protein-driven Mehler-like reaction, and cyclic electron transport. These results suggest that with an oxidised NADPH/NADP^+^ pool, FNR is the preferred route for electrons from Fd_red_. Furthermore, the active YqjM utilises over half of the available electrons released by water oxidation at PSII, increases the electron transport rate through PSI and prevents the transient pooling of electrons in the PETC. Lastly, characterising the cell’s response to electron sink engineering is an important precondition to the rational optimisation of photosynthesis and production strains. We propose that when a strong NAD(P)H-dependent heterologous enzyme is taking electrons directly from the PETC, focusing on improving the overall efficiency of PETC, the enzyme’s activity or localisation are superior strategies to the deletion of AET pathways.

## Acknowledgements

This work was supported by the Academy of Finland (AlgaLEAF, project no. 322754, to YA; Revisiting Photosynthesis, project no. 315119, to YA), the Novo Nordisk Foundation (PhotoCat, project no. NNF20OC0064371, to YA), and the EU FET Open project FuturoLEAF (grant agreement No. 899576, to YA). All the experiments were performed within the PHOTOSYN infrastructure at the University of Turku. Linda Nevala is thanked for technical assistance with immunoblotting.

## Competing interests

The authors declare no competing interests.

## Author contributions

YA conceived the study. MH, LN and YA designed the research. MH and LN performed the experiments. MH, LTW and LN analysed the data. MH, LTW, LN and YA interpreted the results and formed conclusions. LMY and RK provided the custom-made substrate and Syn::YqjM strain. YA secured funding. MH drafted the manuscript, and all authors finalised and approved it.

## Data availability

The datasets analysed during the current study are available from the corresponding author upon reasonable request.

## Notes

### Competing Interest Statement

The authors have declared no competing interest.

