## Supplement Information for "Strong heterologous electron sink outcompetes alternative electron transport pathways in photosynthesis"

### Supplementary Information

**Strong heterologous electron sink outcompetes alternative electron transport and elucidates coordination of electron distribution from Photosystem I**

Michal Hubáček[1], Laura T. Wey[1], Robert Kourist[2], Lenny Malihan-Yap[2], Lauri Nikkanen[1], and Yagut Allahverdiyeva[1]

[1] Molecular Plant Biology, Department of Life Technologies, University of Turku, Turku, 20014, Finland

[2] Institute of Molecular Biotechnology, NAWI Graz, BioTechMed, Graz University of Technology, Graz, 8010, Austria

* corresponding author: Yagut Allahverdiyeva

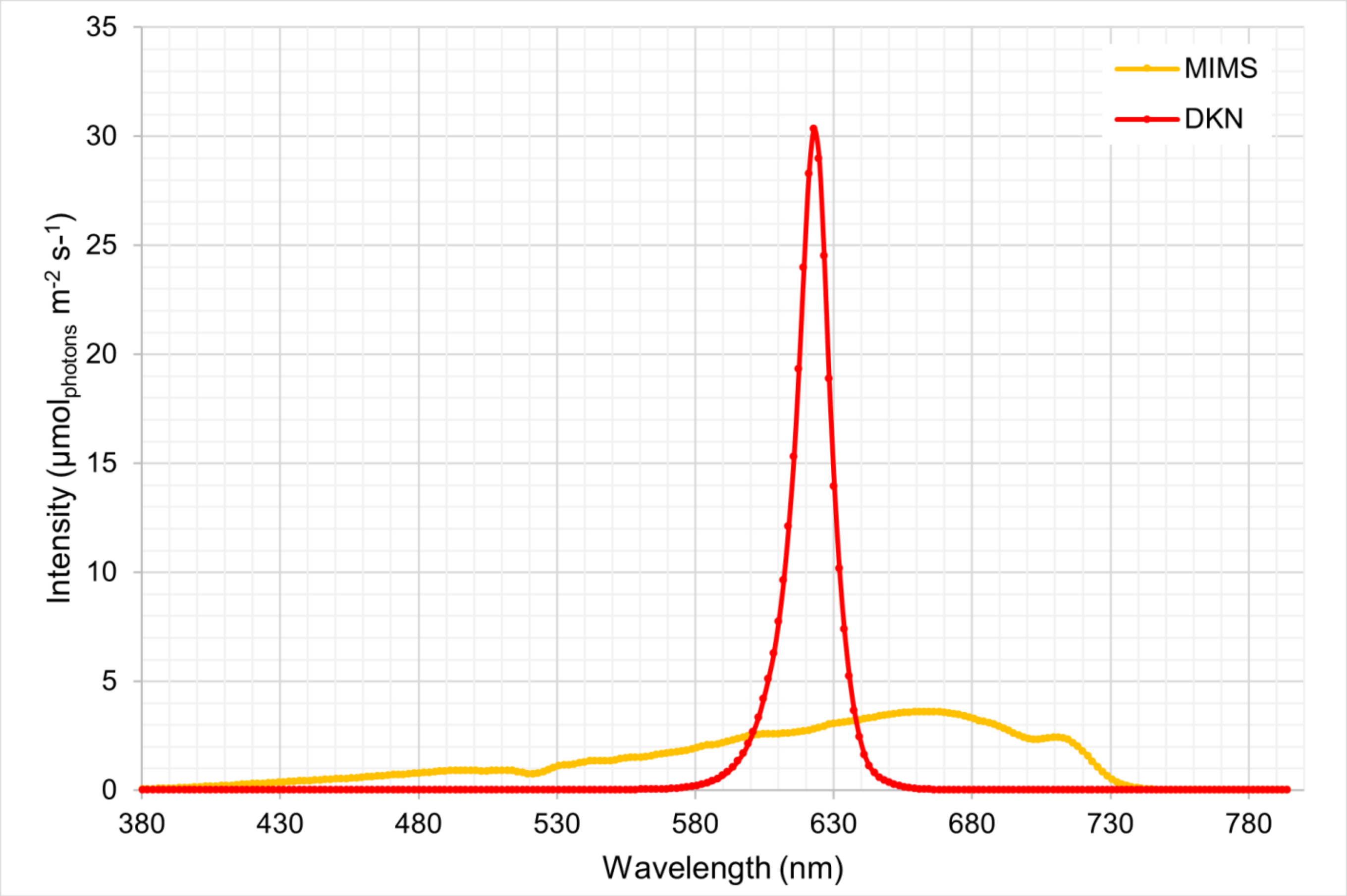


Figure S1. Spectra of the actinic light used in MIMS (broad white light) and DKN (623 nm red light) experiments at PAR at 500 µmol$_{photons}$ m$^{-2}$ s$^{-1}$. Red - DKN, orange - MIMS

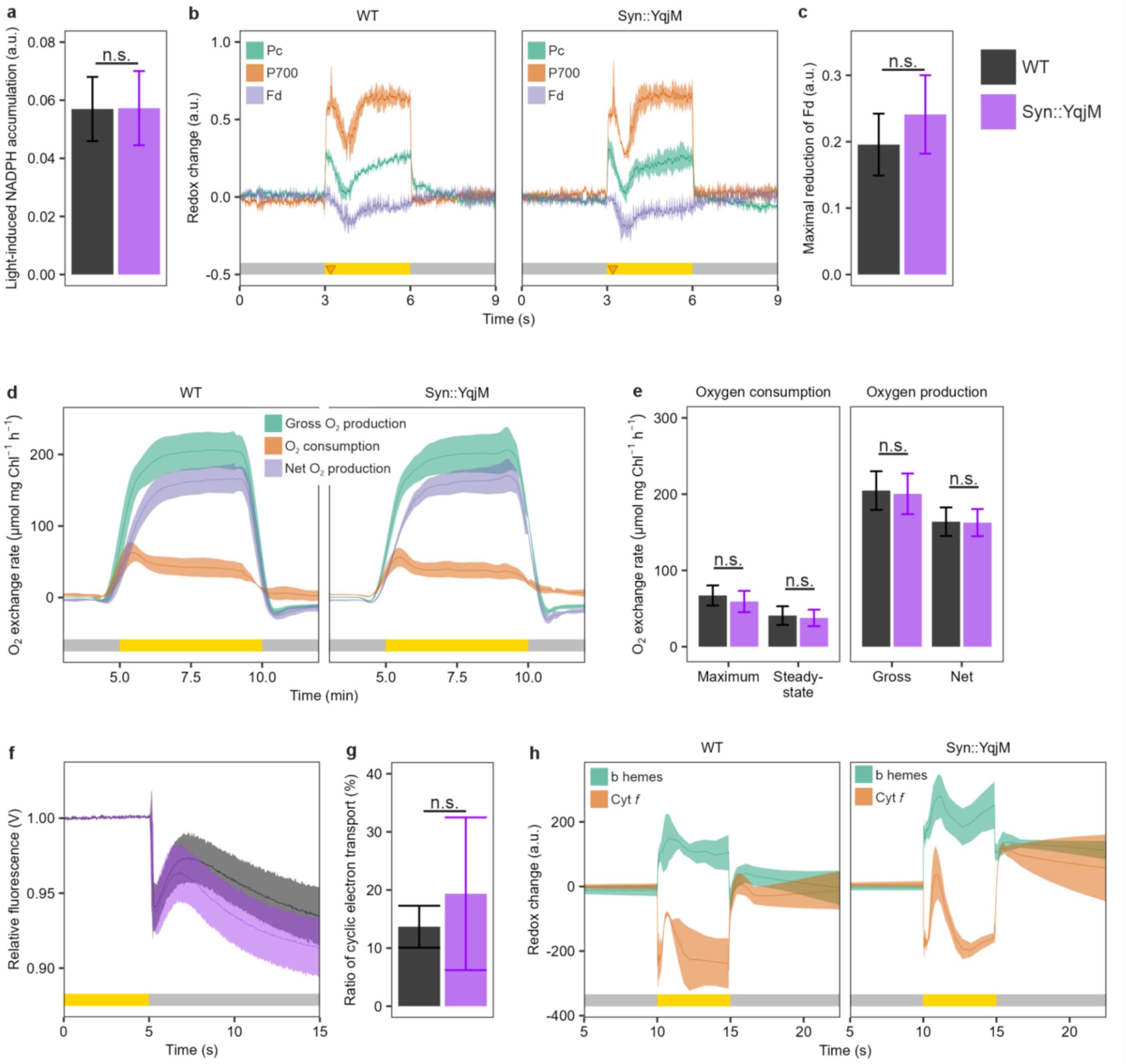


Figure S2. Effect of the expression of YqjM in Synechocystis (WT and Syn::YqjM). **a:** Light-induced NADPH accumulation. **b:** The redox kinetics of Pc, P700 and Fd upon illumination with red actinic light. The orange triangle signifies a saturation pulse distributed to the sample. **c:** The maximal reduction of Fd reached after the saturating pulse was distributed at 200 ms after illumination. **d:** The $O_2$ exchange rate upon illumination with gross and net $O_2$ evolution (green and purple, respectively) and $O_2$ photoreduction (orange). Rates are calculated from a 30s sliding window. **e:** The $O_2$ exchange rate at various stages of illumination. Maximum $O_2$ photoreduction at the peak after dark-light transition and steady-state photoreduction and gross and net evolution were taken as the mean of the $O_2$ exchange rate between 7 - 9 min of measurement. **f:** The post-illumination rise of chlorophyll *a* fluorescence, **g:** The ratio of CET assessed by dark-interval relaxation kinetics of P700 and Pc during illumination. **h:** The redox kinetics of b hemes and Cyt *f* upon illumination with green actinic light. The grey bar in **b**, **d**, **f**, and **h** denotes dark, the yellow bar denotes illumination. Not statistically significant by one-way ANOVA (n.s. > 0.05). Data presented as the mean of 3 biological replicates (4 for Syn::YqjM in **a** and **g**), error bars in column graphs or shading in traces are standard deviations (standard error of the mean in **h**).

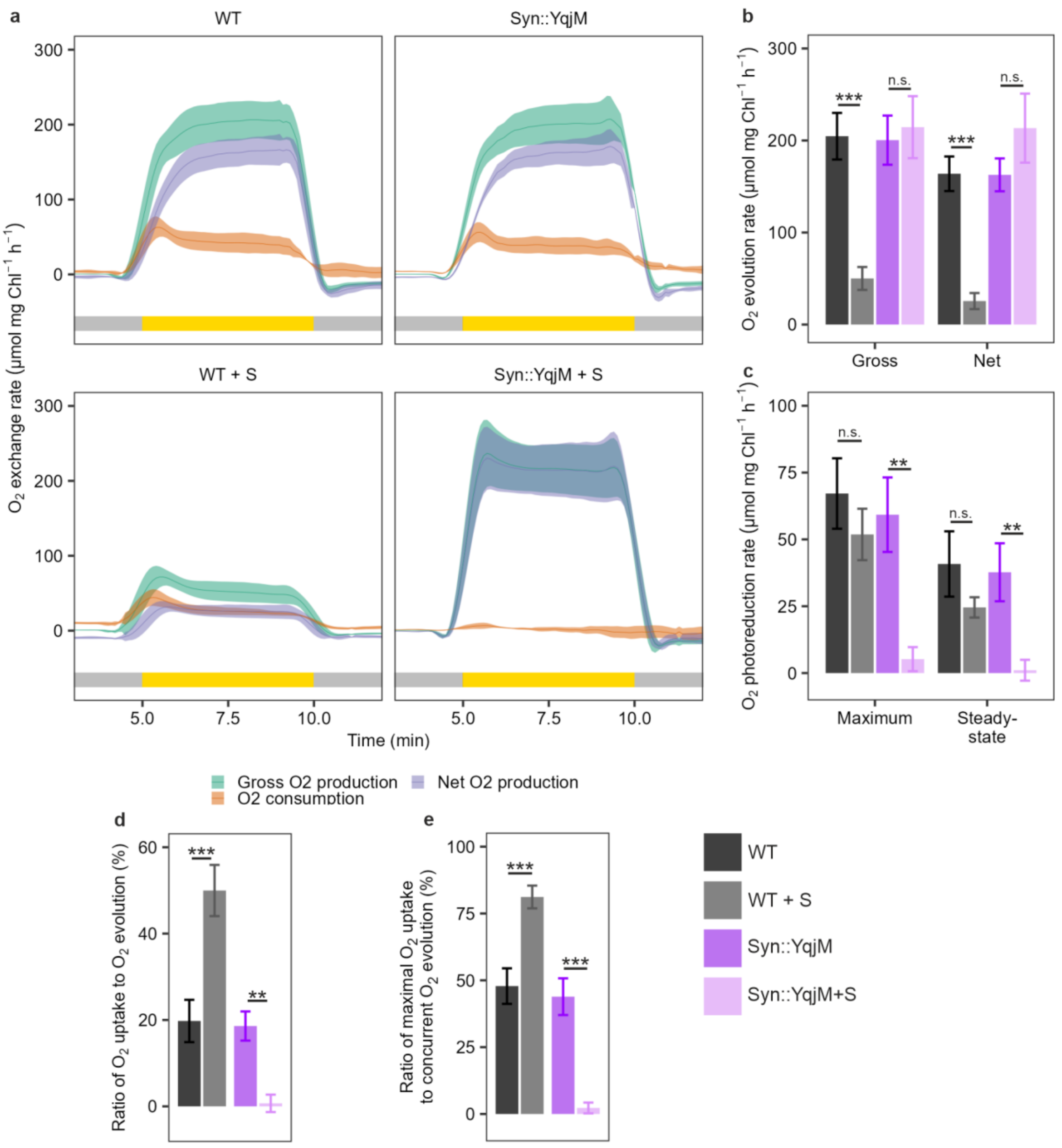


Figure S3. Effect of the substrate on the $O_2$ exchange rate in WT±S and Syn::YqjM±S. **a:** The $O_2$ exchange rate upon illumination. Rates are calculated from a 30s sliding window. **b:** Gross ($p = 3.61 \times 10^{-4}$, $p = 9.05 \times 10^{-1}$) and net ($p = 3.84 \times 10^{-4}$ , $p = 1.04 \times 10^{-1}$) $O_2$ evolution as the mean of the $O_2$ evolution rate between 7 - 9 min of measurement. **c:** Maximum ($p = 3.79 \times 10^{-1}$, $p = 1.40 \times 10^{-3}$) and steady-state ($p = 1.75 \times 10^{-1}$, $p = 3.58 \times 10^{-3}$) $O_2$ photoreduction where steady-state represents the mean of the $O_2$ photoreduction rate between 7 - 9 min of measurement. **d:** The ratio of $O_2$ photoreduction to gross $O_2$ evolution during steady-state ($p = 1.23 \times 10^{-4}$, $p = 4.20 \times 10^{-3}$). **e:** The ratio of maximal $O_2$ photoreduction to the gross $O_2$ evolution at the same time point ($p = 2.73 \times 10^{-4}$, $p = 5.48 \times 10^{-5}$). Data presented as the mean of 3 biological replicates, error bars in column graphs or shading in traces are standard deviations. The coloured bar in **a** denotes illumination: grey - dark, yellow - illumination.

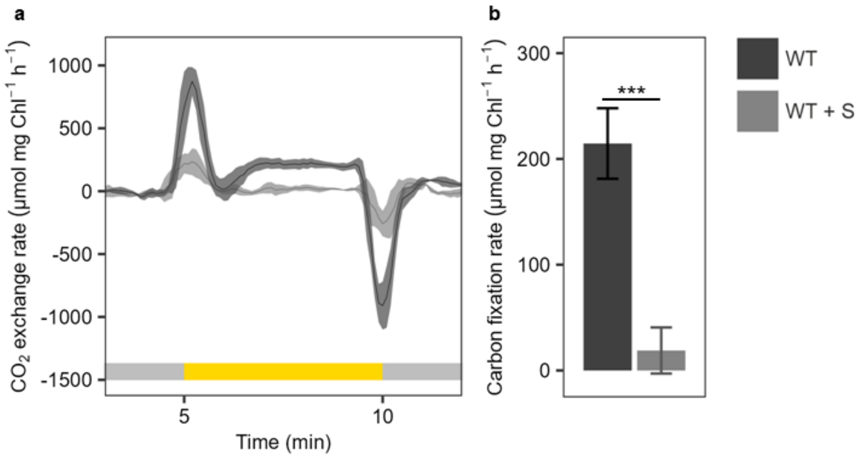


Figure S4. $CO_2$ exchange rate in WT±S. **a:** The $CO_2$ exchange rate changes during illumination. Rates are calculated from a 30s sliding window. **b:** Carbon fixation rate taken as the mean of the $CO_2$ exchange rate between 7 - 9 minutes of measurement ($p = 2.00 \times 10^{-4}$). Data presented as the mean of 3 biological replicates, error bars in column graphs or shading in traces are standard deviations. Statistical significance was tested by one-way ANOVA, n.s. > 0.05, *** ≤ 0.001. The coloured bar denotes illumination: grey - dark, yellow - illumination.

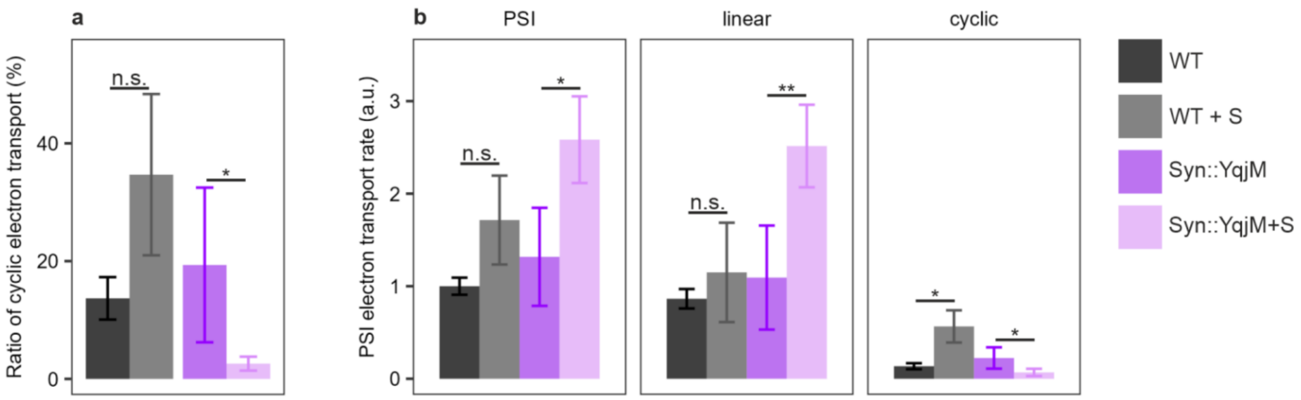


Figure S5. Electron transport through PSI in WT±S and Syn::YqjM±S. **a:** The ratio of CET in WT±S ($p = 6.20 \times 10^{-2}$) and Syn::YqjM±S ($p = 4.40 \times 10^{-2}$). **b:** The electron transport through PSI with WT set to 1 ($p = 6.46 \times 10^{-2}$, $p = 1.16 \times 10^{-2}$) and the contribution of linear ($p = 4.18 \times 10^{-1}$, $p = 7.44 \times 10^{-3}$) and cyclic ($p = 1.37 \times 10^{-2}$, $p = 4.33 \times 10^{-2}$) electron transport. Data presented as the mean of 3 biological replicates (4 for Syn::YqjM±S), error bars in column graphs are standard deviations. Statistical significance was tested by one-way ANOVA, n.s. > 0.05, * ≤ 0.05, ** ≤ 0.01.

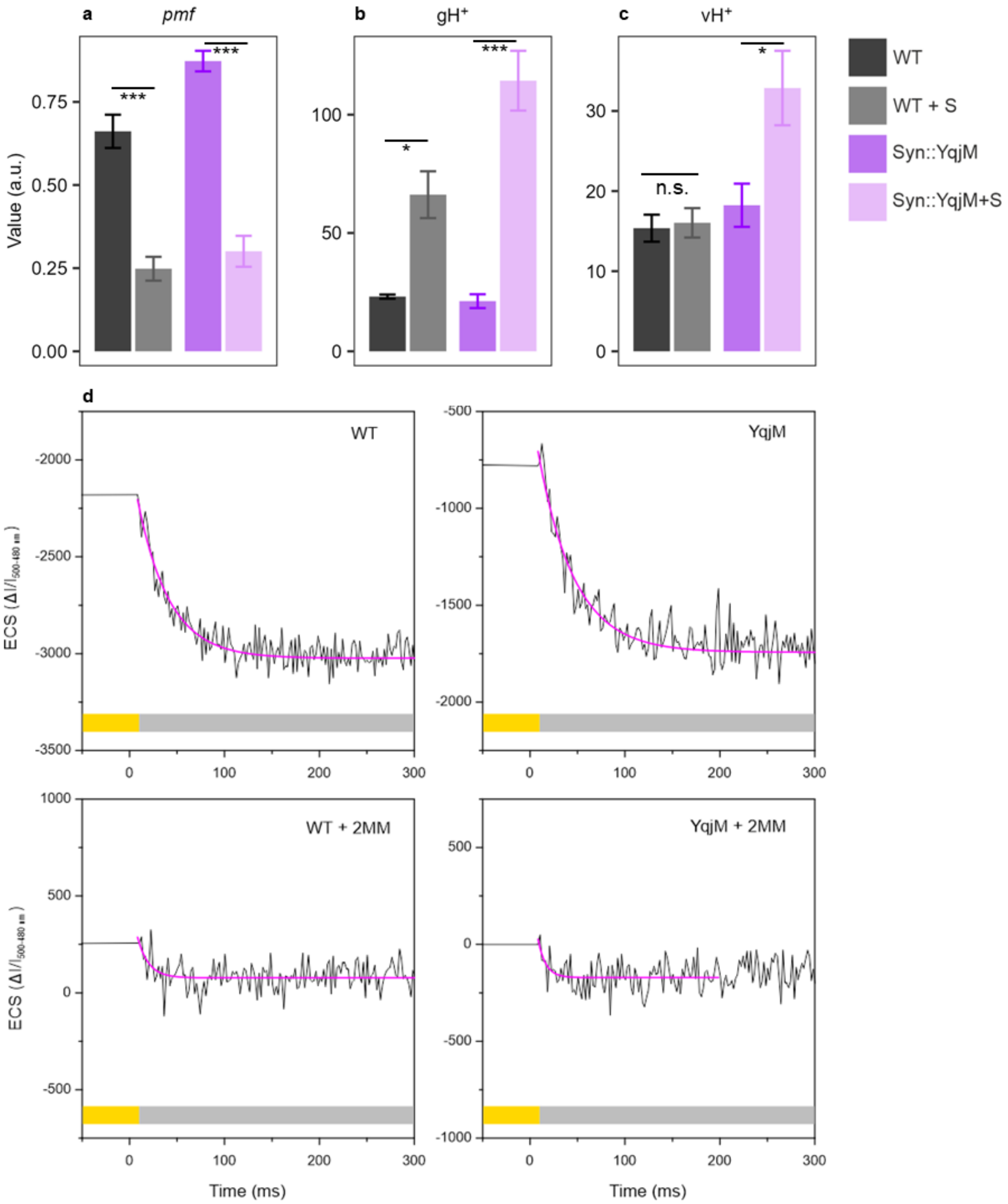


Figure S6. The effect of substrate on pmf generation in WT±S and Syn::YqjM±S. **a:** Proton motive force *pmf* (p = 2.80 × $10^{-4}$, p = 0.00 ). **b:** Thylakoid conductivity $gH^+$ (p = 2.47 × $10^{-2}$, p = 1.40 × $10^{-6}$). **c:** Proton flux $vH^+$ (p = 8.28 × $10^{-1}$, p = 1.84 × $10^{-2}$). Data presented as the mean of 3 timepoints of 1-4 biological replicates, error bars in column graphs are standard errors of the mean. Statistical significance was tested by one-way ANOVA, n.s. > 0.05, * ≤ 0.05, *** ≤ 0.001, **d:** Representative figures of dark interval relaxation kinetics of the ECS signal ($\Delta I/I_{500\text{-}480\ nm}$) for each sample. The purple traces show the first-order fits to the ECS decay kinetics, and the coloured bar denotes illumination: grey - dark, yellow - illumination.

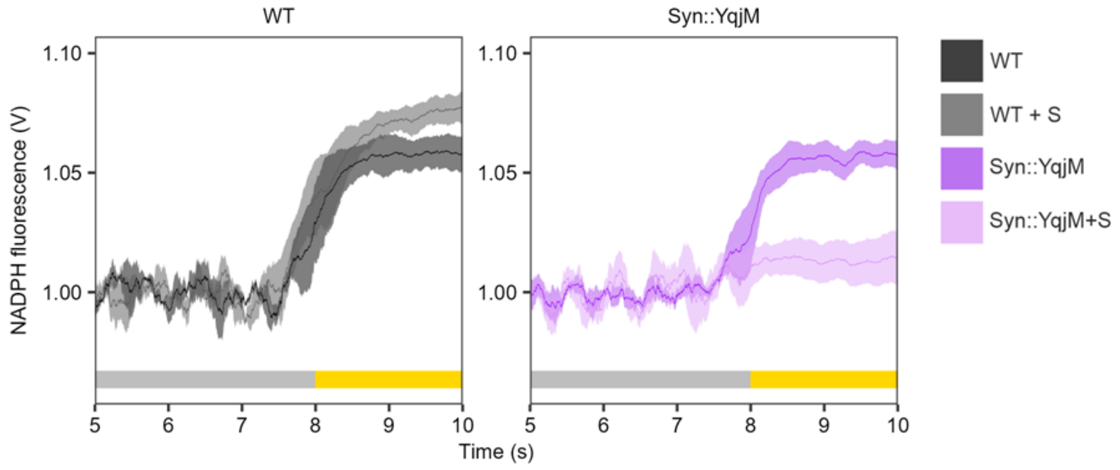


Figure S7. Kinetics of NADPH fluorescence upon illumination by red actinic light. The dark level was arbitrarily set to 1. Data presented as the mean of 3 biological replicates (4 for Syn::YqjM±S), shading in traces are standard deviations. The coloured bar denotes illumination: grey - dark, yellow - illumination.

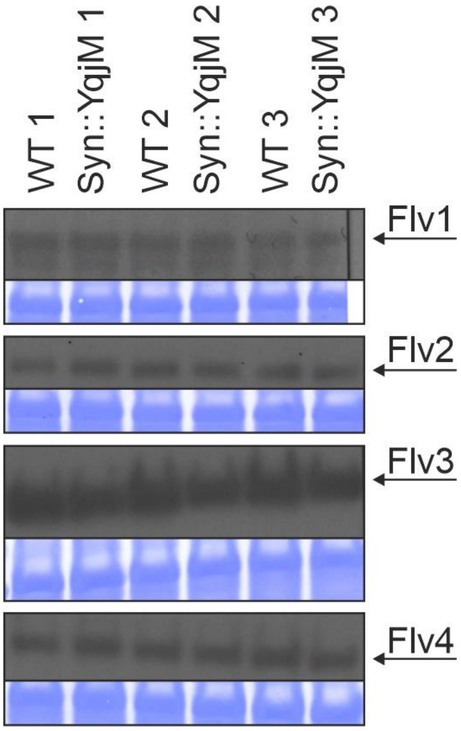


Figure S8. Immunodetection of FDP accumulation levels in WT and Syn::YqjM in the absence of the substrate. Three biological replicates are shown. Coomassie-stained membrane confirmed equal loading between replicates.

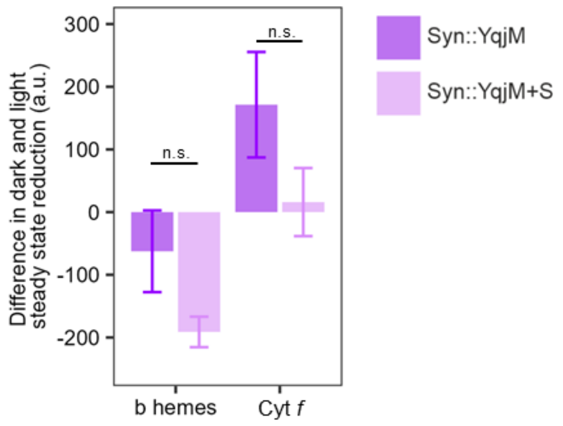


Figure S9. The level of reduction of b hemes ($p = 3.20 \times 10^{-1}$) and Cyt *f* ($p = 3.18 \times 10^{-1}$) after the illumination period in Syn::YqjM±S. Data presented as the mean of 3 biological replicates, error bars in column graphs are standard errors of the mean. Not statistically significant by one-way ANOVA (n.s. > 0.05).

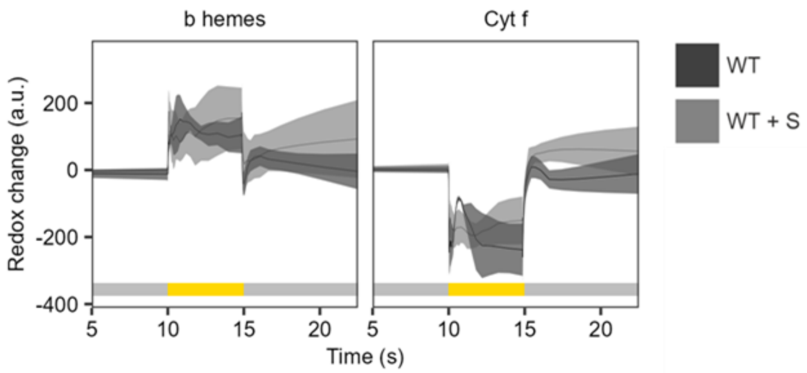

Figure S10. The redox signatures of Cyt $b_6f$ in WT ± S. Data presented as the mean of 3 biological replicates, shading in traces are standard errors of the mean. The coloured bar denotes illumination: grey - dark, yellow - illumination.

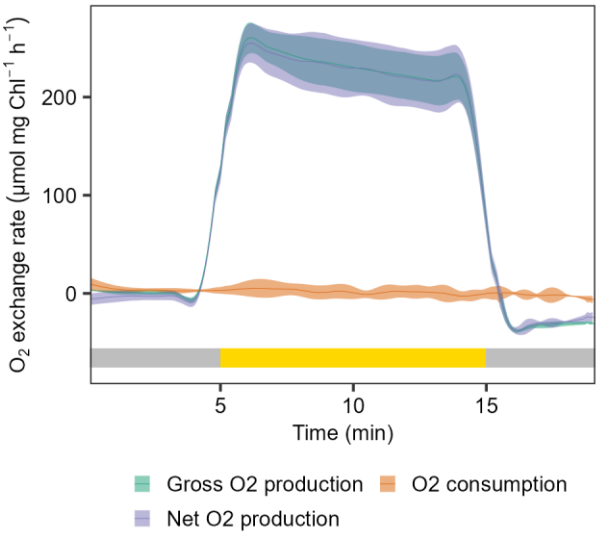


Figure S11. The $O_2$ exchange rate during the biotransformation reaction run in the MIMS sample chamber. Data presented as the mean of 3 biological replicates, shading in traces are standard deviations. The coloured bar denotes illumination: grey - dark, yellow - illumination. Rates are calculated from a 30s sliding window.

### Appendix S1

##### Effect of 2-MM on the thioredoxin and electron transport

2-MM likely impacts the activity of the thioredoxin system. Apart from its impact on the redox regulation of CBB (McFarlane et al., 2019), this alteration could modify the regulation of CET. Even though it has not been investigated in cyanobacteria yet, the activity of the NDH complex has been shown to be affected by the thioredoxin systems in chloroplasts (Courteille et al., 2012; Nikkanen et al., 2018). Additionally, there has been a suggestion for redox regulation of FDPs (Alboresi et al., 2019; Nikkanen et al., 2021). Furthermore, the anchoring of phycobilisomes to the thylakoid membrane may be controlled via thioredoxin-mediated formation and reduction of disulfides between phycobilisome linker peptides (Lindahl and Florencio, 2003). Accordingly, the introduction of N-ethylmaleimide, a thiol-binding molecule similar to 2-MM, results in the inhibition of electron transfer from phycobilisomes to PSI (Glazer et al., 1994). The formation of disulfides in the PsbO protein of the oxygen evolving complex is necessary for PSII activity in chloroplasts (Karamoko et al., 2013), and it is likely crucial in cyanobacteria as well (Guo et al., 2014). The presence of 2-MM may alter this process underlining its inhibitory effect on $O_2$ evolution (Fig. S3).

The addition of the 2-MM substrate to WT cultures led to a sharp decrease in the rate of $O_2$ evolution (Fig. S3a, b), accompanied by an elevation in CET. However, the observed slight increase in PSI electron transport rate compared to untreated cells was not statistically significant (Fig. S5). In Syn::YqjM, steady state $O_2$ evolution was not affected by the presence of 2-MM (Fig. S3a, b), but the PSI electron transport rate showed a significant increase (Fig. S5). These experiments were conducted under strong illumination of 500 $\mu mol_{photons}$ $m^{-2}$ $s^{-1}$. A similar trend was observed in a previous study under weaker irradiance, where the presence of 2-MM reduced PSII yield but did not significantly impact PSI yield (Assil-Companioni et al., 2020). The results obtained suggest that the reduction in PSII activity is balanced by CET, and/or an enhancement in electron influx and a reduction in the outflow of electrons to and from the intersystem chain in the presence of 2-MM.
